# Identification and quantification of modified nucleosides in *Saccharomyces cerevisiae* mRNAs

**DOI:** 10.1101/327585

**Authors:** Mehmet Tardu, Qishan Lin, Kristin S. Koutmou

**Author notes:** **Corresponding author** Kristin Koutmou, 930 N University, Ann Arbor, MI 48105.

## Abstract

Post-transcriptional nucleoside modifications have long been recognized as key modulators of non-coding RNA structure and function. There is an emerging appreciation that the chemical modification of protein-coding messenger RNAs (mRNAs) also plays critical roles in the cell. Although there are over 100 known RNA modifications found in biology only a handful have been identified in mRNAs. We sought to identify and quantify modifications present in the mRNAs of yeast cells using a high throughput ultra-high performance liquid chromatography-tandem mass spectrometry (UHPLC-MS/MS) method that measures the levels of 36 types of RNA nucleosides in parallel. We detected the presence of six modified nucleosides in mRNAs at relatively high abundances: N7-methylguanosine, N6-methyladenosine, 2’-O-methylguanosine, 2’-O-methylcytosine, N4-acetylcytidine and 5-formylcytidine. Additionally, we investigated how the levels of mRNA modifications vary in response to cellular stress. We find that the concentrations of mRNA modifications including N6-methyladenosine and N4-acetylcytidine change in response to heat stress, glucose starvation and/or oxidative stress. This work expands the repertoire of potential chemical modifications in mRNAs, and utilizes a high-throughput approach to search for modifications that highlights the value of integrating mass-spectrometry tools in the mRNA modification discovery and characterization pipeline.

## INTRODUCTION

Cells face the daunting challenge of synthesizing the correct number of proteins at the right time with high fidelity. One way this is accomplished is through chemically modifying proteins and nucleic acids. Modifications change the topologies and chemistries accessible to biomolecules, altering their biogenesis, localization, function, and stability. There have been over 100 RNA modifications identified across phylogeny in non-coding RNAs (Gilbert et al. 2016; Nachtergaele and He 2017). Until recently, post-transcriptional RNA modifications were thought to be largely limited to non-coding RNA species. Over the past five years this dogma has rapidly changed and there is a growing appreciation that a limited set of RNA modifications is also present in protein coding messenger RNAs (mRNAs). Although the list of chemical modifications in mRNAs is growing, only a handful (~15) (Cantara et al. 2011; Machnicka et al. 2013) of RNA chemical modifications have been reported in mRNAs including N7-methylguanosine (m^7^G), N6-methyladenosine (m^6^A), 5-methylcytosine (m^5^C), 5-hydroxymethyl-cytosine (hm^5^C), 3-methylcytosine (m^3^C), 2’-O-Methyl ribonucleosides (Um, Gm, Cm, Am), N1-methyladenosine (m^1^A), and pseudouridine (Ψ) (Carlile et al. 2014; Schwartz et al. 2014; Wang et al. 2015; Delatte et al. 2016; George et al. 2017; Harcourt et al. 2017; Roundtree et al. 2017; Xuan et al. 2017; Bartoli et al. 2018). The discovery of modified nucleosides in mRNAs has garnered a flurry of broad interest because chemical modifications have the potential to modulate every step in the life-cycle of an mRNA following transcription.

Post-transcriptional nucleoside modifications are proposed to alter mRNA stability and folding, protein-recruitment, and translation in a programmed manner (Frye et al. 2016; Gilbert et al. 2016; Roundtree et al. 2017). One modification, N6-methyladenosine (m^6^A), has been reported to impact protein expression by multiple mechanisms, including enhancing mRNA stability (Wang et al. 2014; Shi et al. 2017), cap-independent translation initiation (Zhou et al. 2015) and translation efficiency (Wang et al. 2015; Lin et al. 2016; Li et al. 2017; Shi et al. 2017). Additionally, m^6^A has been linked to a rapidly expanding list of diseases (Hsu et al. 2017; Jonkhout et al. 2017; Angelova et al. 2018), such as obesity (Zhao et al. 2014; Ben-Haim et al. 2015), cervical cancer (Wang et al. 2017), glioblastoma (Cui et al. 2017) and major depressive disorder (Du et al. 2015). Despite the fact that pioneering studies of m^6^A established the capability of mRNA modifications to impact mRNA function with consequences for human health, in general, we still only have a cursory understanding of how mRNA modifications contribute to biology.

Given the extensive diversity of RNA chemical modifications found in non-coding RNA molecules it is possible that the full catalog of modifications present in mRNAs has not yet been uncovered. The current process for discovering and characterizing mRNA modifications is laborious; researchers hypothesize the presence of a particular modification and develop antibody- and/or reverse transcription-based tools to map the position of the modification to the entire transcriptome. While this methodology has yielded multiple ground-breaking findings, it is not a tractable way to fully explore the breadth of possible mRNA modification chemical space. Furthermore, although next generation RNA sequencing (RNA-seq) affords unprecedented accuracy, sensitivity and throughput, the technique indirectly reports on most RNA modifications. The work-flow for these analyses necessarily includes the isolation of an RNA sample (generally via antibody or chemical modification) and analysis of a DNA copy of the isolated RNA, both of which can introduce some biases in sample data. Transcriptome-wide mapping approaches are also limited in their ability to establish absolute nucleoside concentrations. Nanopore sequencing is an emerging technology with the potential to directly detect and count individual modified bases, and while the technique is being rapidly developed it is only useful for mapping modifications that have been identified by other means (Garalde et al. 2018). Thus, high-throughput tools that directly measure mRNA modification levels could provide a useful complement to RNA-seq based approaches by allowing researchers to quantitatively describe the mRNA modification landscape, and providing a means to rapidly and broadly screen potential modifications that may be present in mRNA.

Mass-spectrometry has been used extensively to discover and study protein post-translational modifications, and has the potential to be similarly powerful for directly investigating mRNA post-transcriptional modifications. Furthermore, the technology exists to investigate nucleoside modifications because mass-spectrometry has long been used to examine modifications in the context of non-coding RNAs (Gaston and Limbach 2014; Chen et al. 2018). We therefore sought to quantitatively describe the nucleoside modifications present in yeast mRNAs using a well-characterized ultra-high performance liquid chromatography-tandem mass spectrometry (UHPLC-MS/MS) approach (Deng et al. 2015; Basanta-Sanchez et al. 2016). Our experiments specifically take advantage of a previously published high-throughput variation of UHPLC-MS/MS that uses standards of all 4 unmodified ribo- and deoxyribo-nucleosides, 25 naturally occurring modified nucleosides, and 3 non-naturally occurring modified nucleosides (negative controls) to simultaneously quantify the levels of 36 nucleosides with high accuracy, sensitivity and selectivity, detecting modification levels down to attomolar concentrations (10^−18^ moles/L) (Basanta-Sanchez et al. 2016). Put in other terms, quantitative UHPLC-MS/MS is capable of detecting a single modified nucleoside in one billion yeast cells. Significantly, this method directly detects the presence of modified nucleoside molecules without any potential reverse transcriptase, ligation, or hybridization based-artifacts. We used UHPLC-MS/MS to investigate the nucleoside modification profile of *Sacchromyces cerevisiae* mRNAs purified from yeast grown under non-stressed and stressed conditions (heat shock, glucose starvation and oxidative stress). In our mRNA samples we detected high-levels of four modified nucleosides (m^7^G, m^6^A, Gm, Cm) previously reported in mRNAs, and two modifications formerly thought to be incorporated only into non-coding RNAs (N4-acetylcytidine (ac^4^C), 5-formylcytidine (f^5^C)). The concentrations of all of the modifications we observe in mRNAs change in response to at least one environmental stress. These studies demonstrate how quantitative UHPLC-MS/MS can be used to augment transcriptome-wide mapping approaches in the discovery and investigation of mRNA modifications, and support the supposition that the mRNA post-transcriptional landscape is both complex and dynamic.

## RESULTS AND DISCUSSION

### UHPLC-MS/MS can be used to identify and quantify mRNA post-transcriptional modifications

The extreme sensitivity of UHPLC-MS/MS enables us to detect and quantify every modified nucleoside present in our samples. Consequently, the limit of detection for identifying new mRNA modifications is determined by the purity of our mRNA samples. Therefore, we tested four different purification schemes for yeast mRNAs: single oligo-dT bead pull down, two oligo-dT bead pull-downs, single oligo-dT bead pull-down followed by a RiboZero rRNA depletion kit, and two oligo-dT pull downs followed by RiboZero rRNA depletion. We found the most effective protocol to be composed of two orthogonal steps (oligo-dT pull-down followed by a RiboZero kit), which yielded mRNAs of equal or greater purity than all other tested methods, and at a sufficient concentration for analysis (**Supplemental Figure S1**). Each of our reported values reflects data collected from experiments performed with two biological replicates and three technical replicates of each biological sample.

The key question that we asked regarding this approach was whether the UHPLC-MS/MS signal specific to modifications in mRNAs can be resolved from the noise. Our results indicate that this is indeed possible for abundant mRNA modifications. We approached this problem in three ways: first, by estimating the purity of our mRNA by independent techniques (BioAnalyzer and qRT-PCR); second, by agnostic examination of the distribution of nucleoside modification levels; and third, by inspecting the behavior of known mRNA and non-coding RNA modifications in our UHPLC-MS/MS assay. Independent Bioanalyzer and qRT-PCR assays indicate that our mRNA samples are depleted of rRNAs (5S rRNA, 18S rRNA, 25S rRNA) and a diverse set of tRNAs (tRNA^Arg,UCU^, tRNA^Glu,UUC^, tRNA^Ser,UGA^) (**Supplemental Figures S1 and S2)**.

To assess the distribution of modification levels we compared nucleoside retention level of the modifications (retention of modification = 100% x ([modification in mRNA]/[unmodified nucleoside in mRNA]) / ([modification in total RNA]/[unmodified nucleoside in total RNA])); for example:

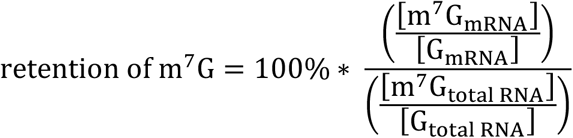

We evaluated the concentrations of each of the 32 modified nucleosides that can be measured with our technique and used these values to estimate the occurrence of modifications per mRNA, for example:

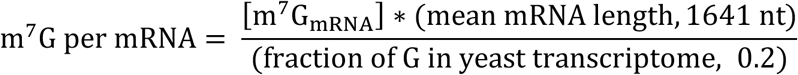

Details of the analysis work-flow are presented in **Supplemental Figure S3**. Our distribution analyses reveal that modifications in our mRNA samples fall into two distinct categories; modifications are either highly-retained and abundant (within 10-fold of m^7^G), or poorly retained and/or rare (> 100-fold less than m^7^G) (**Figure 1, Supplemental Figure S4, Supplemental Table 1**).

**Figure 1:**
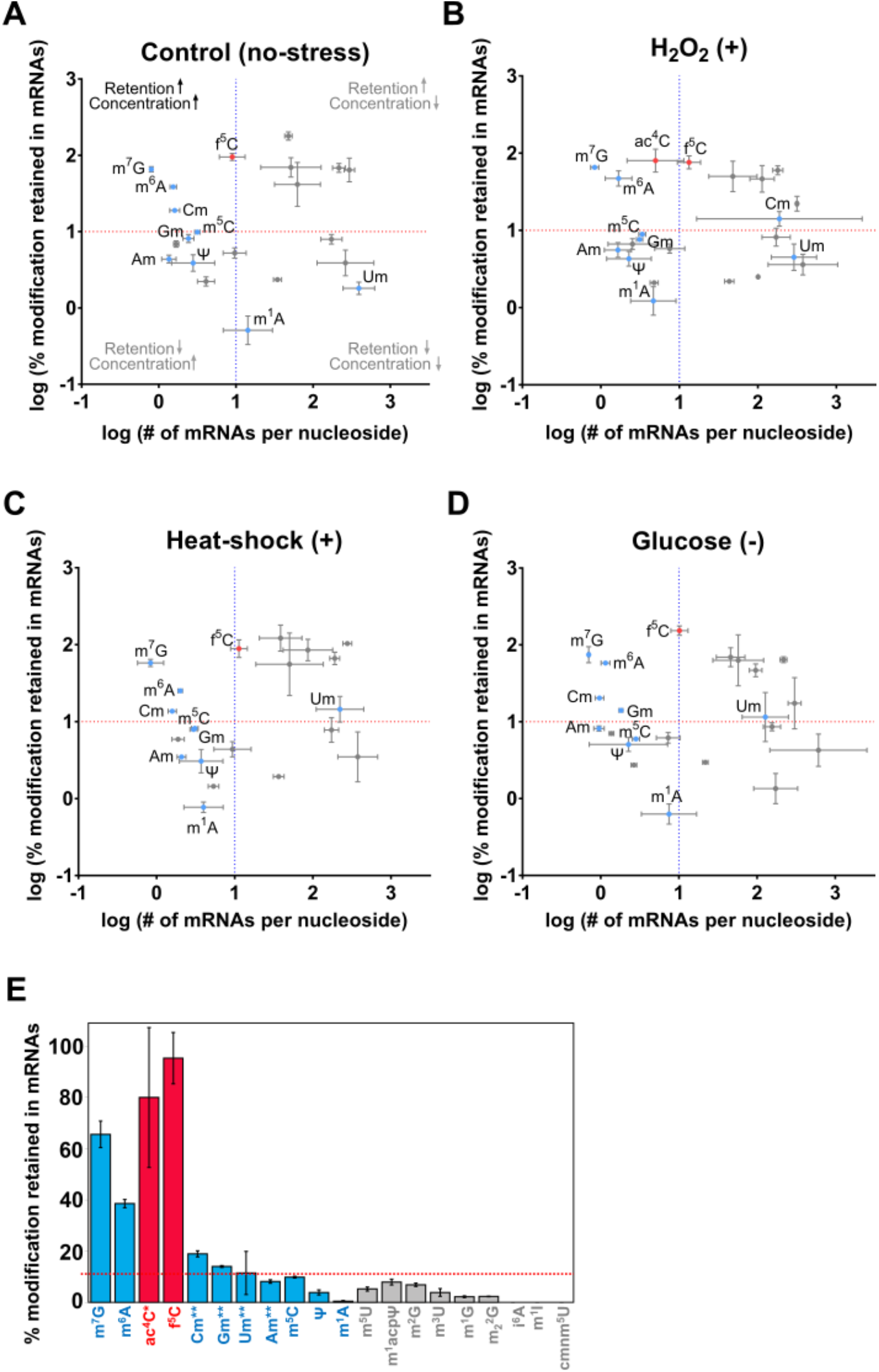
Distribution of modified nucleosides. The retention values (y-axis) and relative concentrations (x-axis) of all modified nucleosides in mRNA samples collected from cells grown under no-stress **(A)**, or stressed **(B-D)** conditions. Nucleosides previously annotated in mRNAs are shown in blue and those that are exclusively in non-coding RNAs are in gray. The values in red represent the two nucleosides that we find in mRNAs that were previously annotated as non-coding species (f^5^C and ac^4^C). **(E)** The percentage of different modifications retained in mRNA samples. Modifications in blue have been previously annotated in mRNAs, and those in gray exclusively in non-coding RNAs. The modifications previously denoted as only in non-coding species (f^5^C and ac^4^C) that we find retained and present at high levels in mRNAs are shown in red.

Next, we identified where known mRNA and non-coding RNA modifications fell in the distributions that we observe. We find that known mRNA modifications are generally well retained at high concentrations, while non-coding RNA modifications typically have negligible retention levels in mRNA samples (**Figure 1**). For example, our results demonstrate that the N7-methylguanosine (m^7^G) modification found on the 5’ end of all mRNAs in eukaryotes is retained our mRNA samples to a high degree (up to 75 ± 18%), and suggest that m^7^G is present approximately once per mRNA (**Figure 1**). We also retain three previously reported mRNA modifications in our mRNAs at high concentrations: N6-methyladenosine (m^6^A), 2’-O-methylguanosine (Gm), 2’-O-methylcytosine (Cm) (**Figure 1E**). In contrast, 90% of modifications previously identified solely in non-coding RNAs are neither retained (< 10% retention in mRNA samples) and/or are not estimated to regularly occur in mRNAs (< 1 per every 100 mRNAs) (**Figures 1 and 2**). For instance, we did not see evidence for the retention of 5-methyluridine (m^5^U, in >95% of tRNAs), 1-methylguanosine (m1G, in ~50% of tRNAs), N2-methylguanosine (m^2^G, in ~60% of tRNAs), N2-N2-dimethylguanosine (m^2^_2_G, in ~60% of tRNAs), N6-isopentyladenosine (i^6^A, in tRNA^Ser^, tRNA^Cys^, tRNA^Tyr^, tRNA^Gly^, (mt)tRNA^Tyr^, (mt)tRNA^Gly^), 5-carboxymethylaminomethyluridine (cmnm^5^U, in tRNA^Leu^ and mitochondrial tRNA^Trp^), 1- methylinosine (m^1^I, found once in tRNA^Ala^), 1-methyl-3-(3-amino-3-carboxypropyl)pseudouridine (m^1^acp^3^Ψ, found twice in 18S rRNA), and 3-methyluridine (m^3^U, present twice in 25S rRNA) (**Figure 1E**). We only observed the retention of two modifications previously identified as non-coding in mRNAs at both levels (80-90%) and concentrations comparable to m^7^G under any experimental condition: 5-formylcytidine (f^5^C) and N4-acetylcytidine (ac^4^C) **(Figures 1, 2, and 3)**. The modifications we observe in our mRNA samples are further scrutinized below.

**Figure 2.**
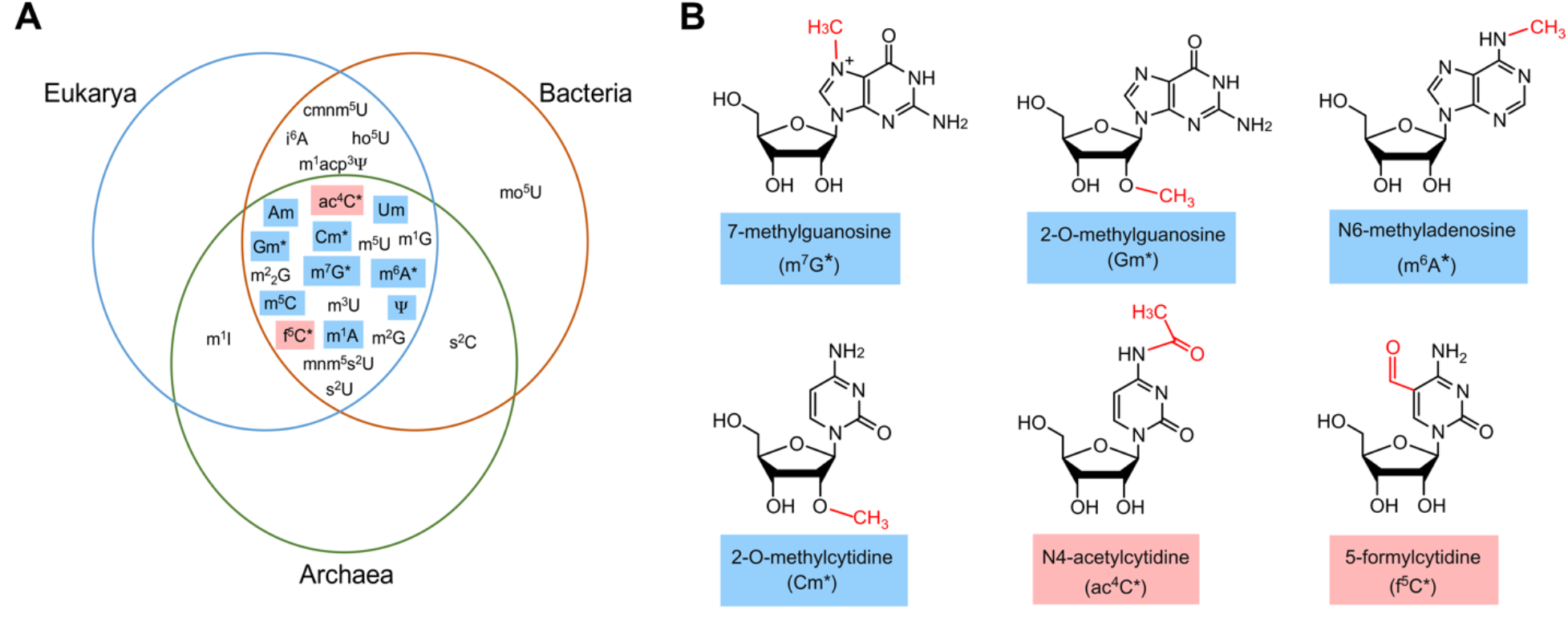
Modified nucleosides measured by UHPLC-MS/MS. **A)** Phylogenetic distribution of the modified nucleosides whose levels we measured in yeast RNAs. Nucleosides previously reported in mRNAs are highlighted in blue. All modifications that we detect above background in mRNAs are denoted with an asterisk (*). Modifications previously observed only in non-coding RNAs are highlighted in red. **B)** Chemical structures of all of the modified nucleosides we observe in mRNAs by UHPLC-MS/MS.

### m^6^A, Cm and Gm are present at high concentrations in mRNA samples

While m^6^A is the most extensively documented mRNA modification we were somewhat surprised to observe appreciable levels of m^6^A in our mRNA samples because the yeast RNA methyltransferase (MIS) complex thought to be solely responsible for m^6^A incorporation is reported to only be active during meiosis (Clancy et al. 2002; Bodi et al. 2015). Although we see m^6^A outside meiosis, the fact that we find m^6^A in yeast mRNAs is consistent with the previous transcriptome-wide mapping study which indicated m^6^A is only found in mRNAs (Bodi et al. 2015). So far, the restriction of m^6^A to meiosis is distinct to yeast. In other organisms, including bacteria, algae, plants, mice, and humans, m^6^A is observed in cells not undergoing meiosis (Fu et al. 2014b; Luo et al. 2014; Meyer and Jaffrey 2014; Chen et al. 2015; Deng et al. 2015). Our results suggest that perhaps yeast is more like other organisms, and can also incorporate m^6^A during additional stages in the cell cycle. Another potential source of m^6^A signal could alternatively be the non-enzymatic Dimroth rearrangement of N1-methyladenosine (m^1^A) mRNA modifications to m^6^A during sample work-up (Macon and Wolfenden 1968). However, Dimroth rearrangement is unlikely to entirely account for the substantial m^6^A signal that we observe (**Figure 1E**).

This left us to ponder the question, if the MIS complex is only expressed during meiosis, where is the m^6^A coming from? We identified at least two possibilities; there may be an additional, not yet identified, methyltransferase that catalyzes m^6^A formation outside of meiosis, or the enzyme that incorporates the N6, N6-dimethyladenosine (m^6^_2_A) modification in 18S rRNA, Dim1, might place a single methyl-group in mRNAs. It has been reported that reduced Dim1 affinity for substrates results in the formation of m^6^A (Desai et al. 2011; Boehringer et al. 2012). If Dim1 binds mRNAs with relatively low affinity this could conceivably provide a mechanism for the non-MIS-complex catalyzed incorporation of m^6^A into mRNAs. We also considered the prospect that our samples were contaminated with rRNAs singly modified by Dim1. However, this seems unlikely because wild-type Dim1 does not generate appreciable levels of singly methylated 18S rRNAs in cells (Desai et al. 2011; Boehringer et al. 2012), and because we find our mRNA samples to be depleted of 18S rRNAs by both qRT-PCR and UHPLC-MS/MS analyses; for example, the m^1^acp^3^Ψ modification found solely in 18S rRNA is not present above background in our samples (**Figure 1**). Our detection of high levels of m^6^A in yeast mRNAs not undergoing meiosis illustrates the value of direct, quantitative approaches for characterizing mRNA modifications.

The conserved methyltransferase Spb1 has recently been shown to incorporate all four 2’O-methyl modified RNA nucleosides into hundreds of mRNAs in yeast (Bartoli et al. 2018). We observe that two 2’ O-methyl modifications (Gm and Cm) are retained in our mRNAs in at least one growth condition; Cm is retained up to 20 ± 0.9%, and Gm is retained up to 14 ± 0.4% (**Figure 1**). We find that 2’ O-methyluridine (Um) is also retained up to 14% in mRNAs, but the high errors (up to 45% error) on the values for this nucleoside do not permit us to annotate it as indisputably present in our mRNA samples (**Figure 1E**). 2’ O-methyladenosine (Am) was retained just under in our 10% samples (up to 8 ± 0.7%). Our data are congruent with the recent discovery that 2’ O-methyl modifications widespread in yeast, and further suggest that 2’ O-methyl modifications are present at high levels in mRNAs.

### m^5^C, Ψ and m^1^A are not retained at high levels in mRNA samples

We did not detect the levels of some other previously identified mRNA modifications above background including Ψ, m^1^A and m^5^C - indicating that our approach may only be able to identify modifications that are abundant in mRNAs relative to other RNA species. This could explain why we find Ψ and m^5^C present just below background in our mRNA samples, but we detect appreciable levels of m^6^A and m^7^G (**Figure 1**). Ψ is estimated to occur in up to 5% mRNA sequences but is highly over-represented in all tRNAs and rRNAs (Carlile et al. 2014), and the prevalence of m^5^C in mRNAs is putatively low but it is found in nearly all cytosolic tRNAs and in 25S rRNA (Machnicka et al. 2013; Legrand et al. 2017). m^1^A is also thought to be relatively rarely incorporated into mRNAs, but is present in over 60% of cytosolic tRNAs and 25S rRNA. It is also possible that m^1^A is not in yeast mRNAs at all, as it has not yet been reported in this organism. In contrast, m^6^A has only been reported in mRNAs in yeast (Bodi et al. 2015), and while m^7^G is both in non-coding and coding RNAs, the preponderance of m^7^G modifications are confined to mRNA species. Thus, while UHPLC-MS/MS is a powerful quantitative approach that can serve as robust screen for mRNA modification discovery, the signal-to-noise ratio in our current implementation only permits us to draw conclusions regarding modifications that are highly-abundant in mRNAs relative to other RNA species.

Our systematic data analyses and exclusive assignment of highly retained, abundant modifications to mRNAs are particularly important in light of the current dialogue regarding prevalence of several mRNA modifications. While the groundbreaking studies that mapped m^5^C and m^1^A to the transcriptome seemed to indicate that these modifications are incorporated at thousands of sites, follow-up reports reach contradictory conclusions - some suggest that these modifications may only be present in a handful of mRNAs, while others support the initial finding that they are common (Edelheit et al. 2013; Li et al. 2016; Legrand et al. 2017; Safra et al. 2017; Guallar et al. 2018; Shen et al. 2018). Thus, despite the limits in the sensitivity of our technique it is still of note that we do not see evidence for high levels of m^1^A in our mRNA samples because this suggests that m^1^A is either not in mRNAs in yeast, or may be incorporated less often than other modifications into mRNAs. Our results neither support nor dispute the ideas that m^5^C is either common or uncommon in yeast mRNAs because while we do not detect m^5^C above background levels in mRNAs, we do see evidence for significant concentrations of the m^5^C oxidation product f^5^C in our mRNAs. These controversies underscore the need for continued development of direct, quantitative methods – such as the UHPLC-MS/MS approach we use here - to detect and characterize mRNA modifications.

### N4-acetylcytidine and 5-formylcytidine are retained at high-levels in mRNA samples

The two nucleosides previously annotated as non-coding chemical modifications that we detected in our mRNA samples, f^5^C and ac^4^C, are conserved in all kingdoms of life, similar to nearly all of the other mRNA modifications identified to date **(Figure 2A, Supplemental Figure S5)**. f^5^C is an *in vivo* oxidation product of the m^5^C through 5-hydroxymethylcytidine (hm^5^C) (Fu et al. 2014a) and has been observed in total RNA from all domains of life and in polyA-enriched RNA fractions from mammalian cells (Huber et al. 2015). Thus, our observation of f^5^C in mRNAs is unsurprising, especially given that hm^5^C has also been reported in eukaryotic mRNAs (Delatte et al. 2016). We verified the presence of f^5^C in our mRNA samples by an additional antibody-based f^5^C detection kit (**Supplemental Figure S6**). Our findings confirm the speculation that f^5^C is in eukaryotic mRNAs (Wang R 2016) and suggest that f^5^C may be quite common. Deoxy-f5C has long been studied in the context of epigenetics (Wu and Zhang 2011) and we expect that many of the tools developed for mapping and quantifying df^5^C, such as aldehyde reactive probes, will prove useful in subsequent studies mapping the location of f^5^C to the mRNA transcriptome.

Ac^4^C is present in both yeast tRNAs and rRNAs and we considered the possibility that our mRNA samples could be enriched for non-coding RNAs containing the modification. However, our qRT PCR studies measuring the levels of rRNA and tRNA species where ac^4^C is found (18S rRNA, tRNA^Ser,UGA^) indicate that these RNAs are depleted to levels far below the level of ac^4^C in mRNAs **(Supplemental Figure S2)**. Additionally, we took advantage of the fact that tRNAs and rRNAs are highly modified molecules to assess our mRNA sample purity in the context of ac^4^C. We reasoned that if tRNA and rRNA fragments containing ac^4^C are present in our mRNA samples, then modifications located in close proximity to ac^4^C on the same tRNA or rRNA should also be retained in our mRNA samples. We evaluated the retention levels of all of the modifications present in all tRNAs where ac^4^C are found **(Figure 3)** and paid particular attention to those modifications in close proximity (2-10 nucleotides) to ac^4^C. We find that modifications in ac^4^C-containing tRNAs are not present above background in our mRNA samples **(Figure 3)**. Additionally, we do not detect the m^1^acp^3^Ψ modification found only in 18S rRNA above background in our samples **(Figure 3)**.

**Figure 3:**
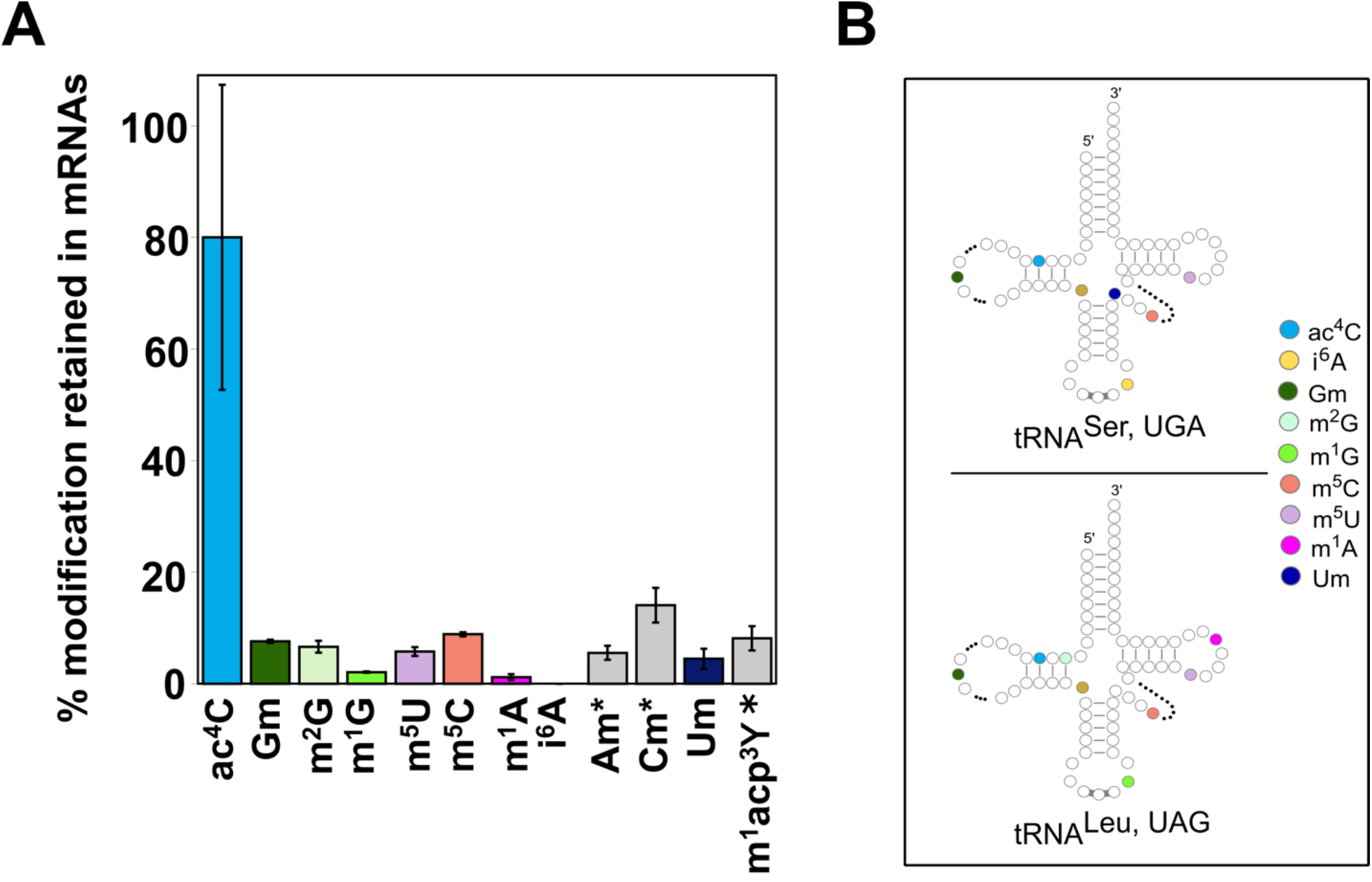
Other modifications in non-coding RNAs where ac^4^C is found are not retained in mRNAs. We plotted the level of modified nucleoside in our mRNA sample relative to the modification concentration present in total RNA. The plot shows the levels of ac^4^C (blue) and modifications on non-coding RNAs that contain ac^4^C retained in our mRNA samples grown under oxidative (H_2_O_2_) stress. **\*** Am, Cm and m^1^acp^3^Ψ are present in 18S rRNA. m^1^acp^3^Ψ is only present in 18S rRNA.

Acetylation is a common biological strategy for reversibly modifying proteins and can change protein-protein and nucleic-acid protein interactions. Non-coding RNAs were already known to contain acetylated nucleosides, and nature appears to have adopted acetylation as a post-transcriptional modification strategy in mRNAs as well. The enzyme that incorporates ac^4^C into mRNAs is not yet identified, but we propose that perhaps ac^4^C could be incorporated into mRNAs by Rra1 (Nat10 in humans), the enzyme responsible for inserting the modification in tRNAs and 18S rRNA (Ito S 2014; Sharma et al. 2015). If this hypothesis proves correct, then identifying the mRNA targets of Rra1 will be of interest because Nat10 is upregulated in a variety of cancers (Shen et al. 2009; Montgomery et al. 2015) and is a potential therapeutic target for premature aging diseases (Balmus et al. 2018). Further work will be required to investigate both the incorporation and modulation of ac^4^C in mRNAs.

### Nucleoside modification levels change in response to cellular stress

One way that mRNA modifications can exert downstream effects could be to rapidly alter mRNA stability and translation in response to the cellular environment. There is precedent for dynamic nucleoside modification in non-coding RNAs in response to environmental stress, nutrition, stages in circadian rhythms, and stage in the cell cycle progression to impact the molecular function (Helm and Alfonzo 2017; Maraia and Arimbasseri 2017). For example, tRNA modifications can mediate the cellular response to stress by controlling the selective, codon-biased translation of particular mRNAs (Maraia and Arimbasseri 2017), and modulations in rRNA modification patterns have been observed in response to disease (Sloan et al. 2017). In mRNAs m^6^A and Ψ incorporation change in response to a variety of conditions such as heat shock, day/night cycles and nutrient deprivation (Carlile et al. 2014; Schwartz et al. 2014; Zhou et al. 2015; Li et al. 2016). The contributions of mRNA modifications to the rapid cellular responses to environmental changes warrants additional careful quantitative characterization. We assessed how mRNA modification levels change in response to stress by measuring the levels of modified nucleosides in mRNA samples collected from yeast grown under three stress-conditions: oxidative stress, heat-shock, and glucose starvation. We find that the levels of mRNA modifications demonstrate statistically significant variations from basal conditions (*p* < 0.005, *p* < 0.01, and *p* < 0.05) in response to environmental challenges **(Figure 4, Supplemental Figure S7 and Supplemental Table 3)**.

The metabolites required for methylation and acetylation (S-adenosylmethionine and acetyl-CoA) change in response to cellular stress. It therefore seems logical that we find that the levels of methylated and acetylated nucleosides changed in response to stress. Our analyses revealed that m^6^A exhibited statistically significant fluctuations in levels (*p* < 0.005) under heat shock (−24 ± 3%), and glucose starvation conditions (+34 ± 10%) **(Figure 4 and Supplemental Table 3)**. This finding is consistent with previous studies reporting that m^6^A levels change in response to heat shock (Zhou et al. 2015). We also saw that the levels of Cm and Gm are increased (+66% ± 12% and 31% ± 9%, respectively) under glucose starvation conditions. In non-coding RNAs 2’ O-methylations are proposed to generally increase non-coding RNA stability (Yang et al. 2015). It is unclear if this is the case in mRNAs. A large portion of 2’ O-methylations in mRNAs are in the region of the molecule that codes for protein (CDS) (Bartoli et al. 2018) where they have been shown to disrupt tRNA decoding by the ribosome (Choi et al. 2018). Normally, disruptions in translation target mRNAs for degradation (Shoemaker and Green 2012). Therefore, it remains to be seen if Gm and Cm moieties in mRNAs will ultimately stabilize or destabilize mRNAs, or if their impact on stabilization will depend on their position in the transcript. Interestingly, we find that N4-acetylcytidine (ac^4^C) is present in total RNA in all of our samples but only detectable in mRNAs purified from yeast grown under oxidative stress **(Figures 1, 3, 4)**. Our results indicate that ac^4^C is one of the most prevalent modified nucleoside species under oxidative stress. This observation, coupled with the conservation of ac^4^C across phylogeny **(Figure 2)** lead us to propose that ac^4^C may have a possible role in regulating the cellular response to oxidative stress in yeast.

**Figure 4:**
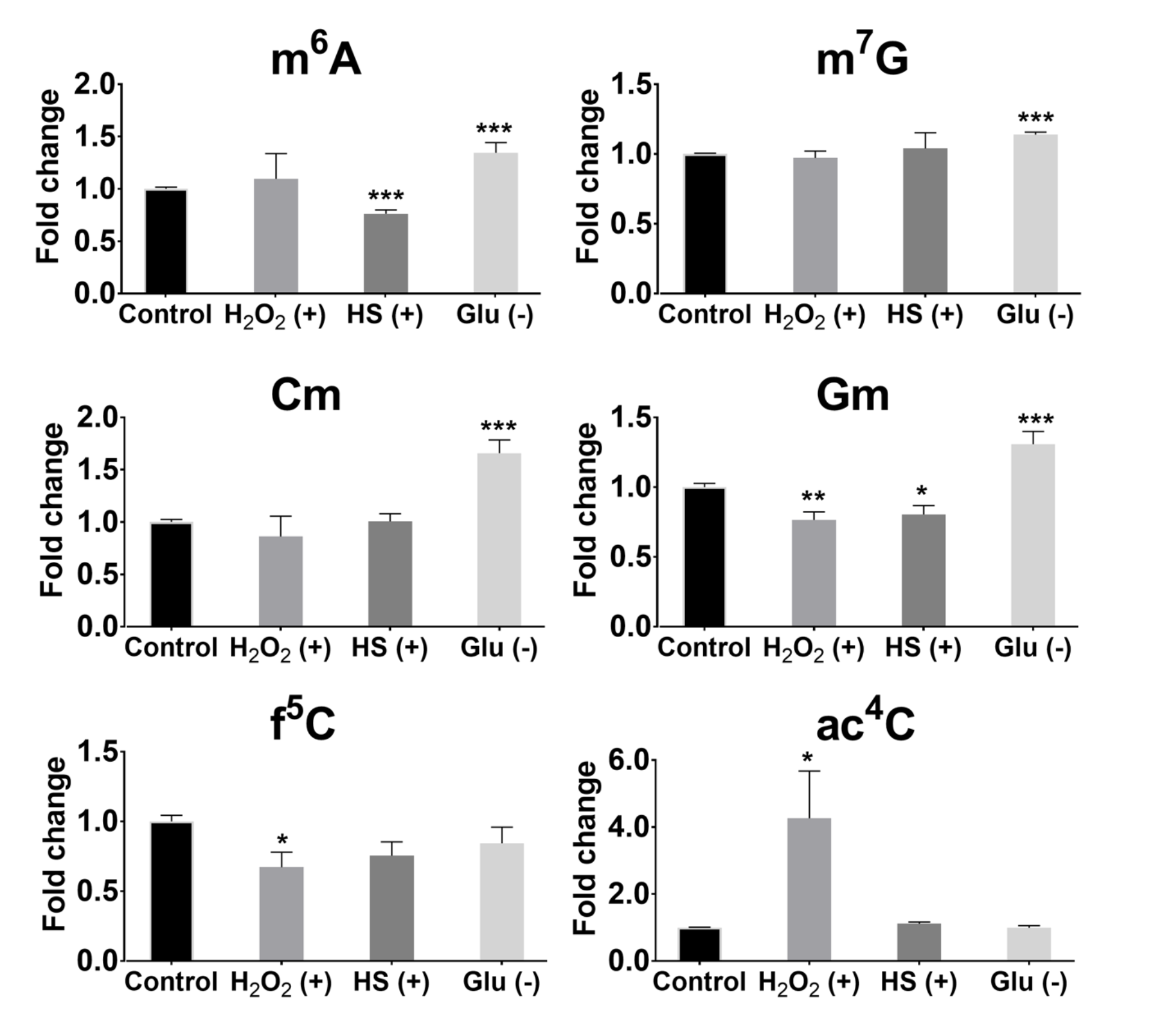
Post-transcriptional modifications in mRNA of *S. cerevisiae* exhibit differential responses to environmental stressors. The fold change for each modification upon stress induction was calculated as the ratio of nucleoside level in the mRNA sample of stress-treated condition ((H_2_O_2_ (+), heat-shock (HS) (+) and glucose deprivation (Glu) (-)) and nucleoside level in the mRNA-enrichment sample of no-stress condition (control). Error bars represent the standard error of mean. \**p*<0.05, \*\**p*<0.01, \*\*\**p*<0.005, Student’s t test.

### Conclusion

The discovery of ac^4^C and f^5^C in yeast mRNAs provides a rational motivation for the development of mapping methods (e.g. based on tools such as those developed in the Meier lab (Sinclair et al. 2017; Thomas et al. 2018)) and transcriptome-wide localization studies of these modifications. More broadly, our findings demonstrate the power of quantitative UHPLC-MS/MS as a tool for mRNA modification discovery. Future improvements in mRNA enrichment techniques will improve the signal-to-noise ratio and enable the detection of less-abundant mRNA modifications, which in turn will allow the field to focus its efforts on mapping and understanding the biology of those modifications. Direct methods such as UHPLC-MS/MS also complement mapping approaches in the study of the biological role of modifications by allowing sensitive and precise quantitation of modification levels. Indeed, our data show that modification levels change in specific ways in response to stress, and thus support the idea that mRNA modification may provide a means for cellular adaptive mechanisms as a rapid response to environmental stressors.

## MATERIALS AND METHODS

### Growth conditions

*Saccharomyces cerevisiae* (BY4741) cells were grown in YPD medium (non-stressed control, oxidative stress and heat-shock conditions) or in defined synthetic complete medium (SC) with 2% glucose (glucose starvation). For all studies, 10 mL of appropriate media (YPD or SC + glucose) was inoculated with individual yeast colony and grown overnight at 30 °C with agitation (200 rpm). Cells were then diluted to OD_600_ = 0.05 into 100 mL of YPD or SC + glucose media and grown to an OD_600_ of 0.6. This culture was subsequently used for stress experiments and sample collection.

Before exposing cells to different stress conditions, 10 mL of cells grown in YPD medium (OD_600_ = 0.6) were collected and used as a control (un-stressed) to compare with stress-induced samples. To assess the effects of oxidative stress on the mRNA modification profiles of *S. cerevisiae*, cells (OD_600_ = 0.6) were incubated with 0.25 mM H_2_O_2_ for 30 minutes at 30 °C. For heat-shock experiments cultures of exponentially growing yeast (OD_600_ =0.6) in YPD medium at 30 °C were heat-shocked by adding an equal volume of fresh medium at 44°C, to immediately reach a final temperature of 37°C. Heat-shocked cells were incubated at 37°C for 45 minutes. Glucose-starvation experiments were carried out by growing cells to OD_600_=0.6 in SC + glucose medium. Then, cells were harvested at 5000 x *g* for 2 minutes, washed three times with SC-glucose medium. After that cells were diluted into fresh SC-glucose medium to OD_600_=0.6 and incubated at 30 °C for 60 minutes. qRT-PCR was performed to measure the mRNA levels of *CCT1*, *HSP30* and *HXT2* genes at different time points for H_2_O_2_ (+), heat-shock (+) and glucose (−) conditions, respectively to verify that stress was induced under each condition (**Supplemental Figure S8**).

### Total RNA extraction, mRNA enrichment and qRT-PCR

Total RNA was extracted from 10 mL of yeast cells (OD_600_ = 0.6) using a standard hot acidic phenol method (Schmitt et al. 1990). Total RNA samples were treated with RNase-free DNase I (Thermo Scientific, USA) (1 U/μg). mRNA isolation was performed in two sequential steps. In the first step oligo-dT magnetic beads (Dynabeads, Invitrogen, USA) were used to selectively bind poly-adenylated RNAs; these beads hybridize to the poly(A) sequence terminating the 3’ end of eukaryotic mRNAs. In the second step, we used a commercial rRNA depletion kit (RiboZero Gold, Illumina) to remove the residual 5S, 5.8S, 18S and 28S rRNAs from our samples. The purity of the isolated mRNA was evaluated using Bioanalyzer RNA 6000 Pico Kit (Agilent, USA) prior to UHPLC-MS/MS analysis. For each sample, rRNA contamination percentage was calculated with the Bioanalyzer software. Additionally, we performed qRT-PCR to measure the levels of tRNA^Arg,UCU^, tRNA^Glu,UUC^, tRNA^Ser,UGA^, 5S rRNA, 18S rRNA, and 25S rRNA to evaluate the purity of our isolated mRNAs.

The qRT-PCRs were performed with Luminaris HiGreen qRT-PCR Master Mix (Thermo Scientific, USA) on StepOnePlus Real-Time PCR System (Applied Biosystems, USA) using gene-specific primers (**Supplemental Table 4**), with *ACT1* (YFL039C) as the internal reference gene. qRT-PCR data was analyzed using the Livak method (2^-(ΔΔCt)^ method). Briefly, average Ct values for all the target genes and housekeeping gene (*ACT1*) in total RNA and mRNA samples were calculated. Then, ∆Ct = Ct (gene of interest) – Ct (housekeeping gene) was calculated for each sample. After that, ∆∆Ct = ∆Ct (mRNA sample) – ∆Ct (Total RNA sample) was calculated. Finally, relative gene level fold change was found by taking 2 to the power of negative ∆∆Ct (Relative gene level fold change = 2^-(∆∆Ct)^).

### UHPLC-MS/MS assay

RNA samples (100ng/10μL each) were analyzed as previously described (Basanta-Sanchez et al. 2016). We used a Waters XEVO TQ-S triple quadrupole tandem mass spectrometry instrument with sensitivity down to 23.01 femtograms, 64.09 attomoles, in approximate closeness of the sensitivity required to analyze and characterize RNA modifications at the single molecular level. Moreover, our samples were separated by a high resolution UHPLC prior to being detected by tandem mass spectrometry instrument, thus further enhancing the selectivity and sensitivity.

Briefly, RNAs were first hydrolyzed to the composite mononucleosides via a two-step enzymatic hydrolysis (nuclease P1 and alkaline phosphatase). Tandem MS analysis of RNA nucleosides was performed on a Waters XEVO TQ-STM (Waters, USA) triple quadrupole mass spectrometer equipped with an electrospray ionization (ESI) source maintained at 500 °C and the capillary voltage was set at 1.5 kV with extraction cone of 14 V. Nitrogen flow was maintained at 1,000 L/h and desolvation temperature at 500 °C. The cone gas flow was set to 150 L/h and nebulizer pressure to 7 bar. Each individual nucleoside modification was characterized by single infusion in positive mode ionization over an m/z range of 100-500 amu. Further nucleoside characterization was produced by using Waters software part of Intellistart MS/MS method development where a ramp of collision and cone voltages is applied to find optimal collision energy parameters for all possible product ions. At least two multiple reaction monitoring (MRM) transitions (precursor/product pairs, one for quantifier and the other for qualifier) were used to monitor each modified nucleoside. To ensure accuracy and make sure at least 12 data point across a LC peak, a scheduled MRM method was developed. The scheduled MRM retention time window of each nucleoside was determined through LC by authentic nucleoside compound (examples of extracted ion chromatograms of ac^4^C and f^5^C given in **Supplemental Figure S5**). To quantify RNA modified nucleosides, calibration curves were generated for 42 modified nucleosides including adenosine, cytidine, guanosine and uridine. [^13^C][^15^N]-G (1 pg/μL) was used as an internal standard. Ion ratio (ratio of qualifier ion/quantifier ion) was used to monitor assay interference. The ion ratio in the RNA digested samples should not change by +/-20% from that of the mean ratio of the nucleoside standards.

A method to extract peak areas from raw data to allow quantification was developed in house using a combination of instrument manufactures suites, MassLynx V4.1 and TargetLynx (Waters, USA). These methods allowed extraction of information to produce calibration curves from each RNA modification standard. In addition, these programs were used to extract the peak areas to be extrapolated on the standard calibration curves for quantification of RNA modifications (quantifications given in **Supplemental Table 1)**. Python script / production of calibration curves as well as quantification from samples was produced in Originlab software suite 2017. All of the raw data are available in **Supplemental Methods**.

### Data analysis

The general work-flow for our assays and data analyses are outlined in **Supplemental Figure S3**. To normalize each of our samples for comparison we first internally normalized the levels of each nucleoside by dividing its molar concentration to the total value of its corresponding unmodified nucleoside molar concentration (e.g. m^7^G_normalized_ = [m^7^G]/([G]]), **Supplemental Table 5**). We next determined the retention value for each nucleoside, as described in main text (retention of modification = 100% x ([modification in mRNA]/[unmodified nucleoside in mRNA]) / ([modification in total RNA]/[unmodified nucleoside in total RNA])) (**Figures 1, 3** and **Supplemental Table 2**).

We estimated the number times each modified nucleoside could be expected to be incorporated per each mRNA transcript. These values are only intended to be used to compare the relative concentrations of modified nucleosides; we are aware that these estimates do not reflect the exact number of times a particular modification is incorporated. To make our estimations, we used the internally normalized the levels of each nucleoside, and multiplied it by the average mRNA length and frequency of each nucleotide in yeast genome to calculate frequency of nucleoside modification per mRNA molecule (e.g. frequency of m^7^G nucleoside = (m^7^G_normalized_) x (average mRNA length) x (frequency of guanine nucleotide in yeast transcriptome). The estimated average mRNA length that we used was 1641 nucleotides (1641 nucleotides = average ORF length (1385 nt) + average total UTRs length (256 nt)) (Hurowitz and Brown 2003). The number of mRNAs per nucleoside is expressed as 1/frequency of nucleoside (**Supplemental Table 2**).

The analysis of the distributions of nucleoside modification retention levels was performed as in (Trang et al. 2015) (**Supplemental Figure S4**). Briefly, the percent retention data (**Supplemental Table 2**) for all experimental conditions were used to identify a cut-off value from the distribution of nucleoside modification retention levels. For this analysis, we used cutoff tool (Trang et al. 2015) to distinguish nucleoside modifications on mRNA than on ncRNA. This tool uses finite mixture models to model bimodal data and to estimate a cut-off value that separates the two unimodal peaks for a given type-I error.

To establish how modification levels vary in response to cellular stress we calculated the fold change of each nucleoside under different stress conditions. This was accomplished by dividing the level of each nucleoside in stress exposed mRNA-enriched sample by the level of the nucleoside in control mRNA-enriched sample (no-stress) (e.g. (m^7^G_normalized,mRNA, stress condition_ / m^7^G_normalized,mRNA, no-stress control_)) (**Figure 4** and **Supplemental Figures S3 and S4**).

### MethylFlash 5-formylcytosine (5-fC) quantification kit to measure f^5^C-levels

The f^5^C level of total RNA and purified mRNA were measured using MethylFlash 5-formylcytosine (5-fC) Quantification Kit (Epigentek, USA) according to the manufacturer’s instructions, including the negative and positive controls. 100 ng of RNA in 10 μl RNase-free dH_2_O was used to estimate f^5^C concentration in the samples according to the standard curve constructed from the manufacturer provided controls (**Supplemental Figure S6**).

## Author Contributions

Mehmet Tardu designed, performed and analyzed all RNA isolation experiments, analyzed the mass-spectrometry data, and wrote the manuscript. Qishin Lin performed the UHPLC/mass-spectrometry experiments, analyzed data, and wrote the manuscript. Kristin Koutmou designed experiments, analyzed data, and wrote the manuscript.

## Notes

The authors declare no competing financial interests.

## Acknowledgements

We would like to acknowledge Dr. Daniel Eyler for his helpful discussion of data analysis. We thank Dr. Markos Koutmos, Dr. Ryan Bailey and Dr. Anna Mapp for their thoughtful reading of the paper and comments.

